# Label-free, real-time on-chip sensing of living cancer cell via grating-coupled surface plasmon resonance

**DOI:** 10.1101/684936

**Authors:** Giulia Borile, Stefano Rossi, Andrea Filippi, Enrico Gazzola, Pietro Capaldo, Claudia Tregnago, Martina Pigazzi, Filippo Romanato

**Author notes:** contributed equally (Stefano Rossi, Andrea Filippi).

## Abstract

The application of nanotechnologies to address biomedical questions is a key strategy for innovation in biomedical research. Among others, a key point consists in the availability of nanotechnologies for monitoring cellular processes in a real-time and label-free approach. Here, we focused on a grating-coupled Surface Plasmon Resonance (GC-SPR) sensor exploiting phase interrogation. This sensor can be integrated in a microfluidic chamber that ensures cell viability and avoids cell stress. We report the calibration of the sensor response as a function of cell number and its application to monitor cell adhesion kinetics as well as cell response to an external stimulus. Our results show that GC-SPR sensors can offer a valuable alternative to prism-coupled or imaging SPR devices, amenable for microfluidic implementation.

## INTRODUCTION

Biomedical research is continuously evolving and the need for more sophisticated and innovative tools is rising mainly due to the contamination of different knowledge.

Cell-based analysis constitutes a fundamental step in life science research, where it is used to assess cell behaviour and response to external stimuli [1]. A gold standard approach to visualize cell response is offered by fluorescence microscopy, which usually requires cell labelling and post-detection analysis, this latter causing the complete loss of information about the kinetic of the process [2]. Moreover, cell adhesion to specific substrates is of increasing interest in the field of 3D culture models aiming to recapitulate *in vivo* complexity with the ease of *in vitro* culture [3].

Surface Plasmon Resonance (SPR) based sensors are of growing interest because of the promising label and enzyme-free, as well as a real-time detection approach for the biological specimens [4], [5].

SPR refers to the collective oscillation of conduction electrons at the interface between a conductor material (typically gold or silver) and a dielectric (i.e. air or water, but also nucleic acids or proteins) upon light interaction [6]. Nanostructured plasmonic biosensors have been developed to improve miniaturization, integration and multiplexing in a lab-on-a-chip approach, while providing high sensitivity and specificity [7], [8].

Recently, applications of SPR sensors as label-free and real-time approach for dynamic cellular analysis have been reported with promising results [9]–[11]. Although most application are based on SPR imaging and focused on antigen detection, and live cell dynamic reactions are detectable as angle of resonance of the SPR changes. This technique allows monitoring molecular interactions occurring within the first few hundreds of nm over a gold surface since the plasmon field is evanescent [6], however it ensures the precise monitoring of phenomena occurring on the cell membrane (like adhesion or rearrangement), regardless of changes in the intracellular compartments, that are far from the SPR sensitive volume. Moreover, the response to external stimuli, was successfully monitored with SPR system implemented for imaging [12], [13]. To date, several challenges need to be addressed to promote the application of SPR-based label-free cell analyses, such as the maintenance of cell integrity during the experimental process. Most of commercially available instruments use glass slides coated with gold, that should be installed on the prism before analysis with a careful handling process to reduce cell damage [9]. Moreover, some instruments can tolerate in the microfluidic components only particulates under few microns, thus cell-SPR application would cause clogging [9].

We previously developed and optimized a nanostructured sensing platform based on grating-coupling and phase-interrogation [14], [15]. This approach was found to be particularly suitable for integration of the plasmonic sensor in microfluidic Polydimethylsiloxane (PDMS) circuits, to allow sensing procedures and tightly control volumes, including multiplexing.

Here, we performed an experimental application of our GC-SPR integrated in a PDMS microfluidic chamber to label-free monitor cell adhesion capability and cell-surface interaction while scaling down medium volume and maintaining cell integrity during manipulation. These results pose the basis for further in vitro studies where cells are maintained in three dimensions cultures for recapitulating human physiological conditions, such as bone marrow niche, that currently rely mainly on microscopy-based analysis.

## MATERIALS AND METHODS

### Numerical Simulations

We performed numerical simulations using the commercial software COMSOL Multiphysics® [16]. The design of the structure replicates the physical dimensions of the grating (duty cycle, linewidth and height) and the field is simulated at a fixed azimuth (as done experimentally) at the resonance angle. Both gold and medium are considered bulk. The same simulation was performed using two refractive indexes: 1.33 for water and 1.38 for cells. The color code adopted to visualize the field represents in red the skin depth.

### Plasmonic Grating fabrication and PDMS embedding

Grating fabrication was performed as previously described by our group [15]. Briefly, digital gold gratings were fabricated, choosing a period of 400 nm and a linewidth of 200 nm, corresponding to a duty cycle of 50%. To fabricate the plasmonic gratings a positive photoresist, Microposit S1805®, was chosen for the laser interference lithography (LIL) [17]. LIL optimal dose was found to be 65 mJ/cm^2^, corresponding to exposure times around 10 min. A 24° incidence angle was used to fabricate gratings with a period of 400 nm, according to the optimization provided in [15]. The grating morphology was characterized by Atomic Force Microscopy. Microfluidic chambers were realized in PDMS with soft lithography and covalently bonded on the SPR chip, as previously described [14].

### GC-SPR detection and analysis Setup

For SPR generation and detection, we used a custom-made bench setup based on phase interrogation, previously adopted for other applications [14]. Briefly, an incident laser beam at 633 nm crosses a half-wave plate mounted on a motorized rotation system, before reaching the sensor. Reflected light is collected by a CMOS camera and the entire system is controlled by a custom-made software. The recorded reflectance is a function of the polarization angle and it is therefore fitted with a harmonic function. A shift of the function phase angle is proportional to the refraction index variation [17].

### Cell Culture

SHI-1 cells (purchased from DSMZ, Germany) were maintained in DMEM supplemented with 10% FBS at 37°C and 5% CO_2_. Prior to the experiments, sensors were coated with Fibronectin (20 mg/mL in Hank’s Balances Salts Solution, HBSS) for 2 hours. After two washouts with HBSS, cells were seeded at the desired concentration.

### Cell Viability Staining

SHI-1 cells were let adhere to the surface, then washed twice in PBS to remove non-adherent cells and debris. To identify cells and evaluate their viability, Calcein-AM (ThemoFisher) was used following manufacturer’s instructions.

### Live imaging with 2-photon microscope

For live cell imaging we used a custom-build 2-photon microscope, previously described by our group for different applications [19]. Briefly, excitation wavelength was set at 800 nm and Calcein-AM emission was imaged with a 525/40 nm bandpass filter. Images were processed and analyzed with FIJI software. All data are reported as mean± s.e.m. (standard error of the mean).

## RESULTS AND DISCUSSION

### GC-SPR in conical mounting characterization

In conical mounting (fig. 1a), the reflectance exhibits a sinusoidal dependence as a function of light polarization angle [15] that is acquired and fitted as in fig. 1b. A change of the refractive index over the grating causes a shift in the reflectance spectra (fig. 1b) that was previously characterized and optimized by our group [15].

**Figure 1.**
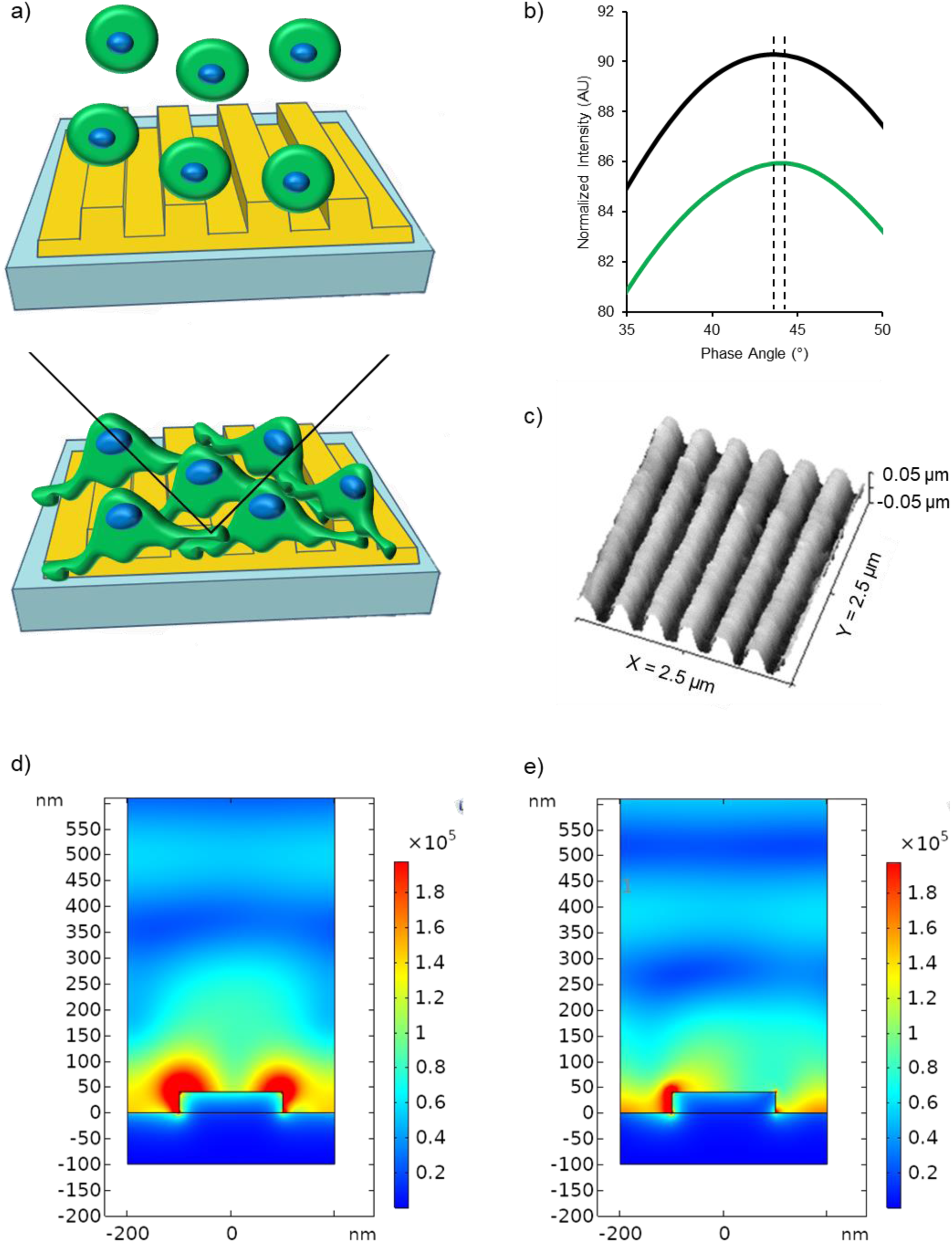
a) Representation of the experimental approach used to interrogate cell adhesion with GC-SPR. Cells in suspension (top) adhere to the gold grating (bottom). Black line indicates the incident light path and the reflected light path used to detect signal. b) Details of polarization spectra acquired in no cell condition (black) and with cells adherent to the surface (green). The dotted lines indicate the spectra maximum, calculated interpolating experimental data to a sinusoidal curve. c) Grating morphology reconstructed by Atomic Force Microscopy. d) COMSOL simulations of the field over the grating in water and (e) in a medium with cell refractive index (1.38).

SPR performance relies on fabrication parameters, that were previously identified in 400 nm period and 40 nm line depth in aqueous medium [15]. Thus, morphological analysis of the fabricated samples was performed by Atomic Force Microscopy (see fig. 1c). The obtained profiles revealed a good agreement between the fabricated grating and the theoretical parameters. All the gratings used for the experiments reported in this paper were tested for fabrication parameters.

Then, we aimed to evaluate the penetration depth of the field into a biological material. To reach this goal we used the COMSOL Multiphysics® software. It allows to design the desired structure, where materials are represented by their complex permittivity, and exploit a Finite Elements Method to solve its interaction with incident light [17]. Since cells are big with respect to the nanostructure scale, they were treated as a bulk. In this way, we could achieve information about the surface plasmon evanescent field shape and penetration in the cell. The skin depth, visualized in red in fig. 1d and 1e, is well below 100 nm in water and even less in a medium with the estimated average refractive index of a cell. This means that the large part of plasmonic field probes the cell plasma membrane, that constitutes the first 10 nm over the adhesion layer [19]. This aspect is crucial, since it allows to monitor changes in the adhesion of cells over the gold surface, independently of rearrangements occurring inside the cell cytosol, that is too far from the sensitive volume. In comparison with optical microscopy techniques, this confinement of the acquired signal to the first layers over the grating makes the system intrinsically “confocal”, and the data obtained belong to an “optical section” of around 1% of the cell height.

### Fibronectin coating as adhesion layer and anti-fouling agent

SHI-1 is a bone marrow (BM)-derived myeloid leukemia cell line that grows in suspension but can adhere to surfaces upon a coating with proteins typical of the extracellular matrix. We identified the fibronectin, that in its soluble form is highly present in blood plasma and is one of the components of the extracellular matrix, suitable for our aims. The most effective adhesion coating resulted in fibronectin at a concentration of 20 mg/mL for 2 hours. We thus monitored with our GC-SPR system the coverage of the sensitive area with fibronectin that reached a steady state after 80 minutes with a sigmoid curve, as expected (fig. 2a, representative data of three different experimental replicates, Hill fit: R^2^=0.98). Interestingly, we observed that fibronectin coating acted as an anti-fouling agent. This effect is important since it avoids any other interfering agent being adsorbed to the grating or involved in any reaction. This is of crucial relevance while doing experiments with viable cells considering that their growth medium is full of salts, proteins and vesicles coming both from the cells themselves and from serum added to culture medium. To assess whether SHI-1 cells could be affected by the gold grating, we stained cells with Calcein-AM, a standard indicator that becomes fluorescent only in viable cells. A representative image of cells loaded with Calcein-AM and acquired at the interface between gold and glass, is shown in fig. 2b. Regardless of the substrate, cells accumulated Calcein-AM and showed fluorescent signal that appears stronger on the gold substrate because of reflection. In three experimental replicates we acquired random fields of fluorescent cells and compared cell area (Gold: 214±8 µm^2^ vs. Glass: 195±7 µm^2^, n=50 cells per condition, T-test not significant, p-value=0.1) that did not show significant differences between the two substrates. Then we wanted to compare cell morphology using standard shape descriptors like Aspect Ratio (Gold: 1.57±0.07, vs. Glass: 1.58±0.08, n=50 cells per condition, T-test not significant, p-value=0.9) and Roundness (Gold: 0.67±0.02, vs. Glass: 0.68±0.03, n=50 cells per condition, T-test not significant, p-value=0.8) and even this analysis did not revealed significant differences. Thus, the gold grating does not affect SHI-1 cells adhesion or viability.

**Figure 2.**
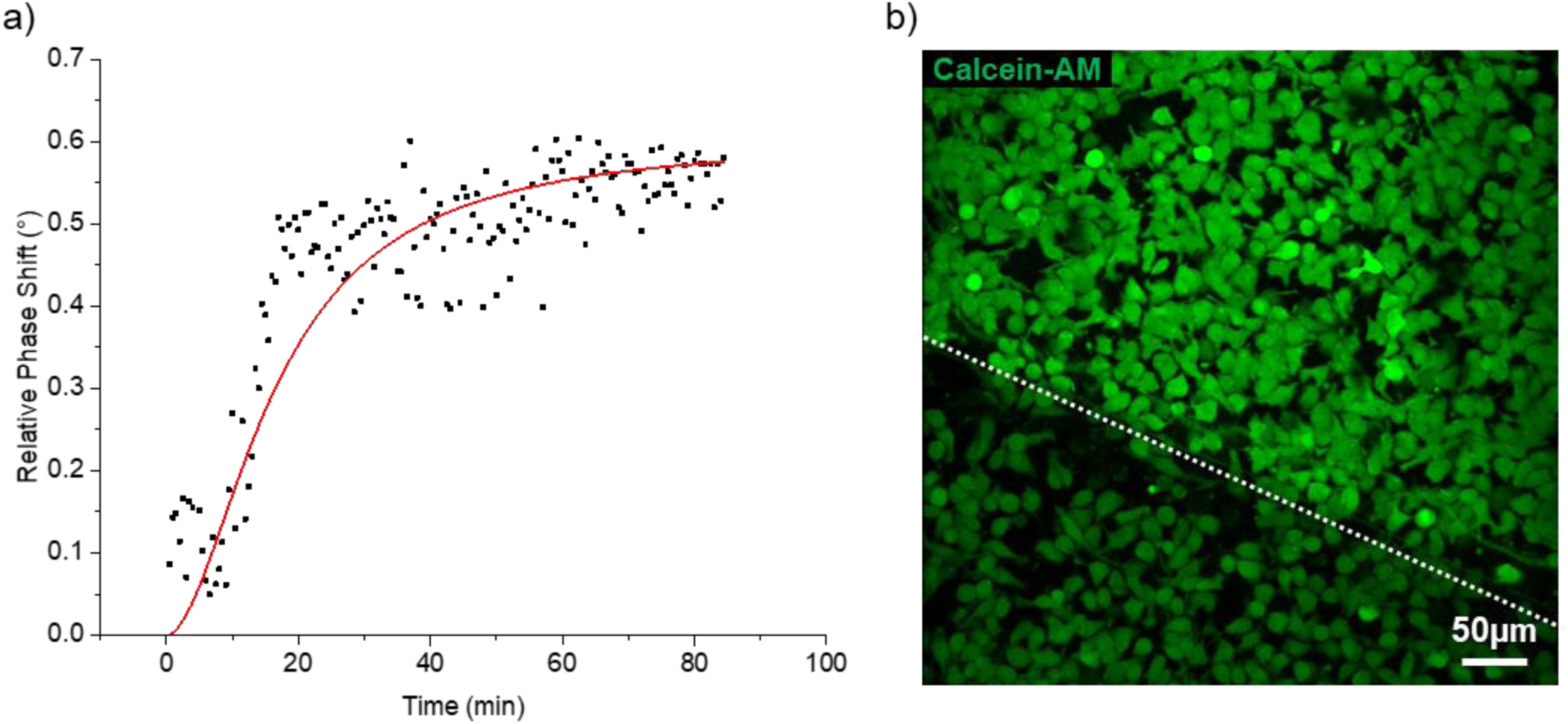
a) Fibronectin coating on gold grating monitored as SPR phase angle variation in the microfluidic chamber. Representative data of one out of three different experimental replicates, in red Hill fit: R^2^=0.98. b) SHI-1 cancer cells after seeding on fibronectin coating, imaged at the interface between the regions where the substrate is either glass (below the dotted line) or gold grating (over the dotted line). Calcein-AM staining of living cells did not highlight differences between the two surfaces, nor differences were observed in the cell shape as quantified in the results section. Area_Gold_: 214±8 µm^2^ vs. Area_Glass_: 195±7 µm^2^; AspectRatio_Gold_: 1.57±0.07, vs. AspectRatio_Glass_: 1.58±0.08; Roundness_Gold_: 0.67±0.02, vs. Roundness_Glass_: 0.68±0.03; n=50 cells per condition, T-test not significant).

### GC-SPR label-free quantification of adherent cells

SHI-1 cells were then used to monitor and quantify cell adhesion to the selected coating. The adhesion of SHI-1 cells over the fibronectin-coated gold grating causes a large shift in the phase angle proportional to the surface coverage, that is correlated to number of adherent cells. Using different cell concentrations, it was possible to build a calibration scale of the SPR response of our system (fig. 3a representative trace from single experiments, fig. 3b mean value and s.e.m. of three experimental replicates). The minimum detectable number of cells (limit of detection) corresponded to around 100 cells/mm^2^, while the maximum angle shift (sensor saturation) was registered for numbers over 2000 cells/mm^2^ (Δα=0.91°±0.02°). This last parameter is in good agreement with a similar experiment where authors reported that saturation was reached with 1600 cells/mm^2^ in a prism-coupled SPR device (ΔR=0.78°) [10]. Thus, our GC-SPR based system results in a global response comparable to that previously obtained in a similar experimental report. This result supports the use of SPR technology towards cell-based assays.

**Figure 3.**
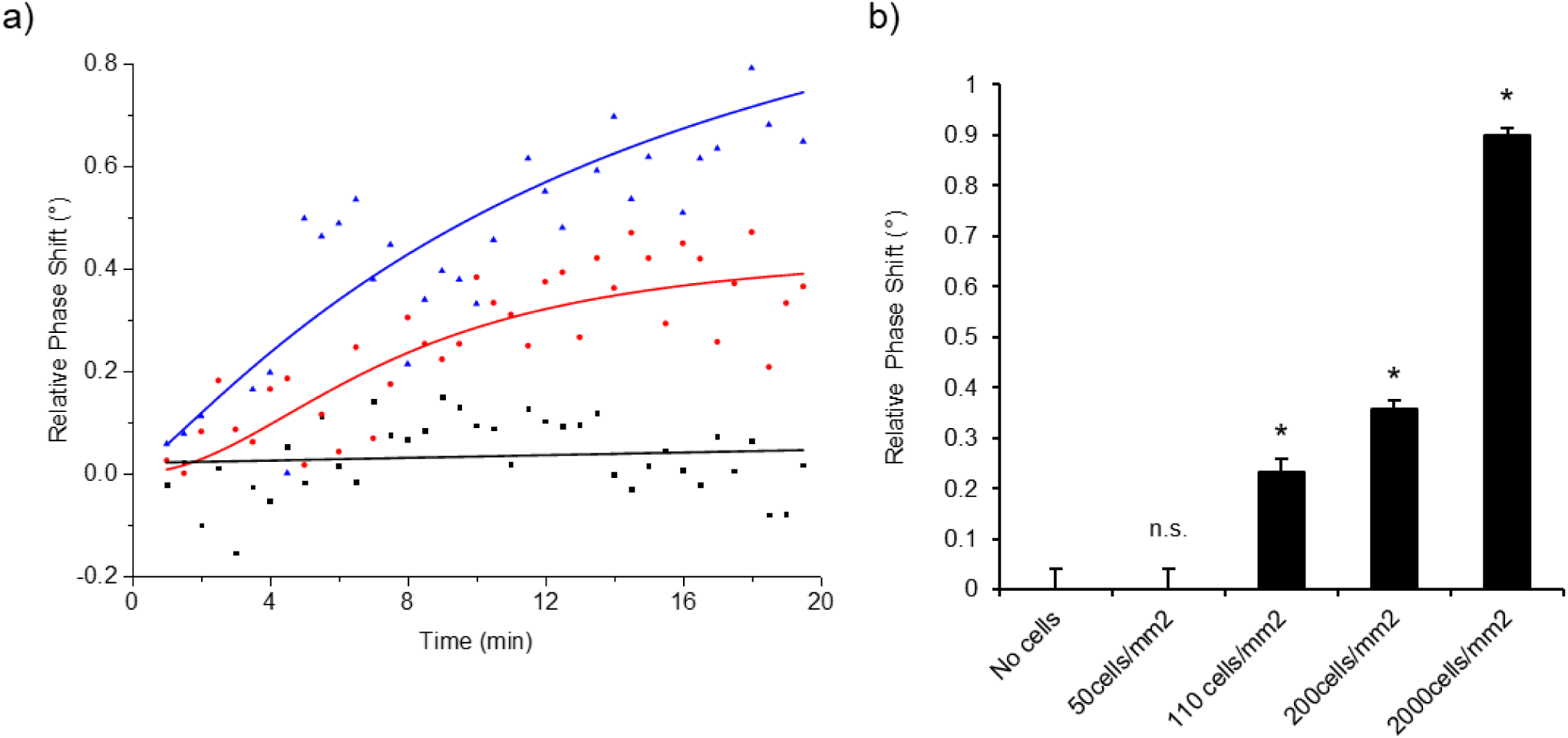
a) SHI-1 cancer cells were monitored during adhesion to coated gratings at room temperature. Black dots represent cell medium signal, red dots represent the final concentration of 200cells/mm^2^ and blue dots 2000cells/mm^2^. SPR phase spectra were acquired every 10 seconds. Phase shift (with respect to baseline) was calculated with a custom MatLab® routine, then fitted with a Hill function. b) Calibration of the SPR response as a function of cell concentration. Histogram bars represent the mean±s.e.m. of at least three experimental replicates. T-test was performed comparing with no cell condition: * means p-value < 0.05, n.s. means not significant.

### GC-SPR label-free real-time monitoring of cell detachment

To further confirm that SPR can detect morphological changes of cells, we challenged adherent cells with trypsin, an enzyme that causes structural shrinking and detachment from the fibronectin coating. The SPR response resulted in a Δα=-0.72° consisting in a reduction of the cell membrane-covered surface around 95%, consistent with the microscopy analysis performed by fluorescence imaging (see fig. 4).

**Figure 4.**
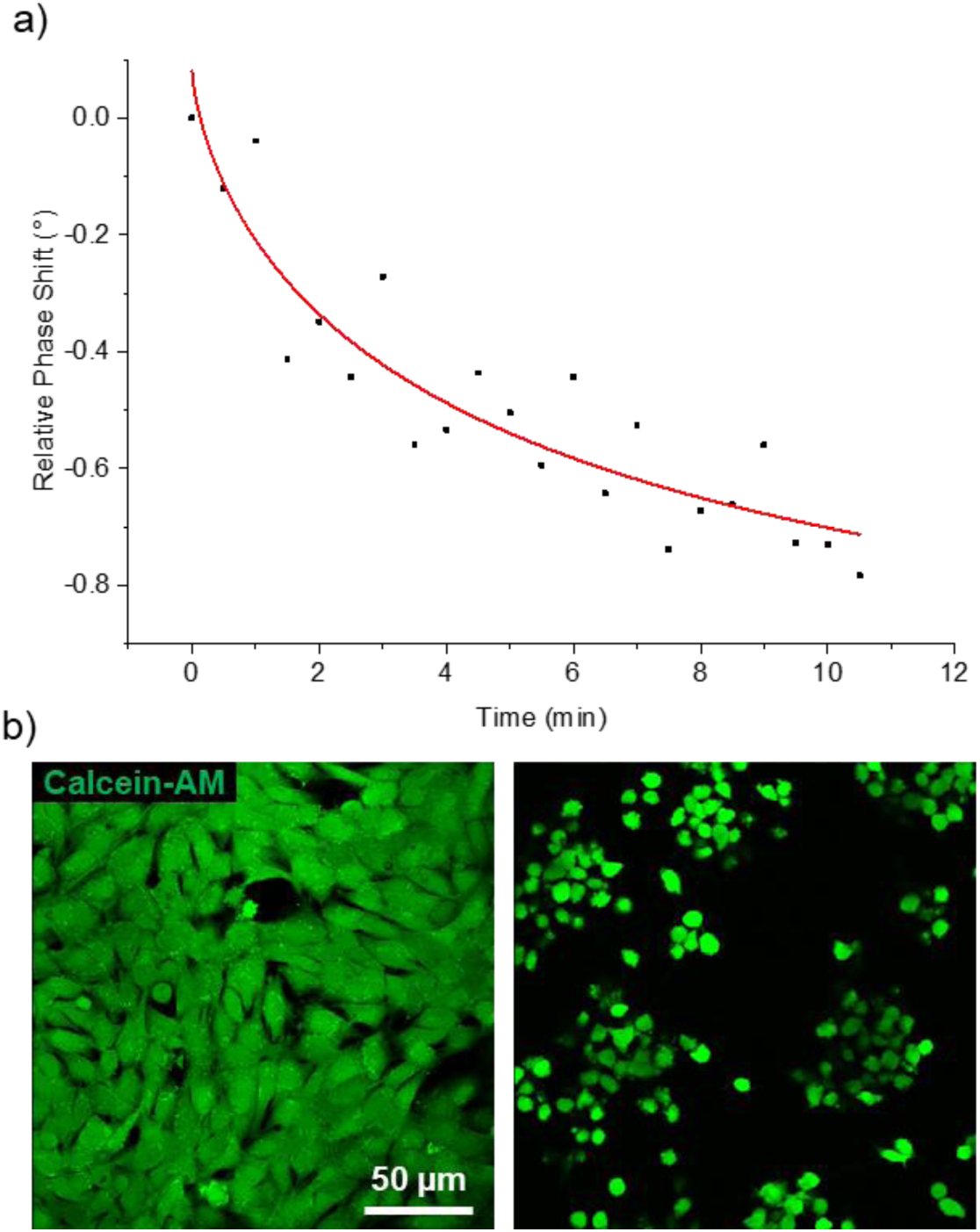
a) SHI-1 cancer cells were treated with trypsin to detach them from grating. As cells detach from the sensing surface, a reduction in SPR signal can be observed. Red curve represents an exponential fit of the data. The curve is representative of three experimental replicates. b) Representative images of cells loaded with Calcein-AM before (left) and after (right) trypsin administration, respectively.

## CONCLUSIONS

In this proof-of-principle paper we have shown that grating-coupled SPR biosensors can be used for live cell dynamics investigation. Our GC-SPR system is based on phase interrogation, that means a sinusoidal behavior of the reflectance as a function of incident light polarization. The SPR signal, in terms of phase shift, is proportional to the surface coverage over the grating, allowing a calibration for cell number quantification. The major advantage of this technique relies on the compatibility with custom made PDMS chambers that allow to seed, maintain in an incubator, and analyse cells without further manipulation, a procedure that is not allowed by standard prism-coupled devices where the gold slide maintained in a petri dish for seeding must be removed and mounted on the system, with the risk of cell damaging. Moreover, we showed that SPR sensors can be used for live cells experiments not only in the imaging configuration. Indeed, in SPR imaging, signal results from amplitude modulation at a fixed angle, unlike GC-SPR where a phase angle scan is performed. Because they are based on intensity interrogation, SPR imaging sensors suffer from worse performance with respect to angle scanning systems ([20] and for a complete review see [21]). In conclusion, our methodological approach results in good resolution, compact, inexpensive and very simple cell detection setup, and opens for further research strategies based on the integration of the sensitive grating into a microfluidic chamber that could improve cell biology studies.

## ACKNOWLEDGEMENT

G.B. acknowledges the support from Fondazione Istituto di Ricerca Pediatrica (Grant: MyFirstIRP). The authors declare no conflict of interest.

## ABBREVIATIONS

SPR: surface plasmon resonance
GC: grating coupled

